# Benchmarking transcriptional deconvolution methods for estimating tissue- and cell type-specific extracellular vesicle abundances

**DOI:** 10.1101/2024.02.27.582268

**Authors:** Jannik Hjortshøj Larsen, Iben Skov Jensen, Per Svenningsen

**Author notes:** Corresponding author: Per Svenningsen. These authors contributed equally to this work (Jannik Hjortshøj Larsen, Iben Skov Jensen).

## Abstract

Extracellular vesicles (EVs) contain cell-derived lipids, proteins, and RNAs; however, the challenge to determine the tissue- and cell type-specific EV abundances in body fluids remains a significant hurdle for our understanding of EV biology. While tissue- and cell type-specific EV abundances can be estimated by matching the EV’s transcriptome to a tissue’s/cell type’s expression signature using deconvolutional methods, a comparative assessment of deconvolution methods’ performance on EV transcriptome data is currently lacking. We benchmarked 11 deconvolution methods using data from 4 cell lines and their EVs, *in silico* mixtures, 118 human plasma, and 88 urine EVs. We identified deconvolution methods that estimated cell type-specific abundances of pure and *in silico* mixed cell line-derived EV samples with high accuracy. Using data from two urine EV cohorts with different EV isolation procedures, four deconvolution methods produced highly similar results. The four methods were also highly concordant in their tissue-specific plasma EV abundance estimates. We identified driving factors for deconvolution accuracy and highlight the importance of implementing biological knowledge in creating the tissue/cell type signature. Overall, our analyses demonstrate that the deconvolution algorithms DWLS and CIBERSORTx produce highly similar and accurate estimates of tissue- and cell type-specific EV abundances in biological fluids.

## Introduction

Body fluids contain a mosaic of cell type-specific extracellular vesicles (EVs); however, establishing which tissues and cell types contribute most EVs is challenging and thus limits our understanding of how EVs contribute to health and diseases. Several animal models have been developed to confidently link cell types and their EVs by genetic reporters (Neckles et al. 2019; Gupta et al. 2020; Luo et al. 2020; Estrada et al. 2022; Hegyesi et al. 2022; Nørgård et al. 2022; Rufino-Ramos et al. 2022); yet, establishing this link in humans is challenging. While traditional methods such as flow cytometry can robustly quantify cell type-specific EVs, flow cytometry relies on a restricted number of cell type-specific protein-based markers that limit its scalability.

Nonetheless, advances in EV RNA analyses have provided new approaches to predict the EVs’ cellular source by computational methods termed transcriptome deconvolution. The deconvolution methods can untangle the tissue- and cell type-specific EV abundances in body fluids; however, their accuracy has not been thoroughly evaluated, and a comparative assessment of deconvolution methods’ performance on EV transcriptome data is currently lacking. Most transcriptome deconvolution methods have been developed to estimate cell type composition of tissues from bulk RNA sequencing. A general premise for these deconvolution methods is that transcript expression from each cell type is linearly additive, enabling estimation of the tissue’s cell type composition from a list of cell type-specific marker transcripts. This reference signature can be established from isolated cells, single-cell, or single-nuclei transcriptome data sets. The marker-based deconvolution methods use this gene list to determine which cell type markers carry the most information for a regression approach to estimate cell type proportions in bulk RNA sequencing data accurately (Newman et al. 2019; Tsoucas et al. 2019; Wang et al. 2019; Jew et al. 2020). In addition to estimating cell type proportions in bulk RNA sequencing data, the deconvolution methods, such as CIBERSORTx (Newman et al. 2019) and Bisque (Jew et al. 2020), have been used to estimate tissue- and cell type-specific EV abundance using plasma and urine EVs (Shi et al. 2020; Zhu et al. 2021; Dwivedi et al. 2023). Moreover, EV-origin has been created for the deconvolution of EV abundances in plasma (Li et al. 2020; Yu et al. 2020). However, transcriptome deconvolution methods make assumptions about the bulk RNA sequencing data and signature matrix’s RNA and cell distribution. For example, deconvolution is most accurate when the bulk and reference transcriptome are from the same cellular compartment, e.g., the whole cell (Sutton et al. 2022). Yet, EVs are considered a separate compartment from the cytosol, and RNA transcripts are not always distributed equally between cells and EVs (Almeida et al. 2022; Padilla et al. 2023). Moreover, the EV secretion rate between cell types is highly variable (Garcia-Martin et al. 2021, 2022; Auber and Svenningsen 2022); thus, rare cell types in the signature may contribute a significant number of EV RNA reads in the bulk samples and *vice versa*. These factors may affect the deconvolution methods’ performance for accurately estimating tissue- and cell type-specific EV abundances, and the factors influencing their results remain to be determined.

A significant hurdle for evaluating the deconvolution methods’ performance in estimating EV abundances is the lack of samples with known proportions of cell type-specific EVs. While EVs isolated from a cell-conditioned medium are useful for assessing performance in pure systems, biological fluids contain a mixture of cell type-specific EVs. The deconvolution methods’ ability to unmix these samples may, therefore, not be sufficiently assessed using only single-cell type samples. An essential fact about transcriptome deconvolution methods is that they, rather than proportions of cell types, estimate the proportion of RNA derived from each cell type (Zaitsev et al. 2019; Sutton et al. 2022). Thus, synthetic mixtures with biological heterogeneity in known proportions can be created *in silico* by sampling and mixing RNA sequencing reads from pure samples (Dong et al. 2023). By leveraging this approach, we used cell and EV transcriptome data from 4 cell lines to create a dataset containing pure EVs, *in silico* mixtures of two cell type-specific EV samples at 5 different proportions (0%, 25%, 50%, 75%, 100%) to evaluate 11 deconvolution methods (Table 1) ability to estimate cell type-specific EV abundance accurately. We tested the effect of adding cell type-specific EV missing in the signature matrix. Finally, we evaluate the concordance in EV estimates of transcriptome deconvolution of plasma and urine EVs and highlight the importance of biological knowledge in establishing tissue/cell type signatures.

**Table 1:** The benchmarked deconvolution methods.

| Algorithm | Regression | Reference |
| --- | --- | --- |
| Bisque | Non-negative least squares | (Jew et al. 2020) |
| CIBERSORTx | Support vector regression | (Newman et al. 2019) |
| EV-origin | Support vector regression | (Li et al. 2020) |
| DeconRNASeq | Non-negative least squares | (Gong and Szustakowski 2013) |
| dtangle | Linear mixing model | (Hunt et al. 2018) |
| DWLS<sup>a</sup> | Dampened weighted least squares | (Tsoucas et al. 2019) |
| DWLSj<sup>a</sup> | Dampened weighted least squares<sup>b</sup> | (Tsoucas et al. 2019) |
| DWLS OLS<sup>a</sup> | Ordinary least squares | (Tsoucas et al. 2019) |
| DWLS SVR<sup>a</sup> | Support vector regression | (Tsoucas et al. 2019) |
| MuSiC | Weighted non-negative least squares | (Wang et al. 2019) |
| MuSiC nnls | Non-negative least squares | (Wang et al. 2019) |

<sup>a</sup> The DWLS-based methods use the same algorithm to identify signature genes for tissue/cell types.
<sup>b</sup> Unlike DWLS, DWLSj does not iterate the proportion estimation to converge on a solution.

## Materials and methods

### Cell line and cell-conditioned medium EV RNA sample data

For the cell lines and their EVs, we used RNA sequencing data from two studies that performed paired analyses of cultured cell lines (Hinger et al. 2018; Almeida et al. 2022). Hinger et al. (2018) used colorectal cancer cells (CRC) and their EVs, and Almeida et al. (2022) used prostate cancer cell lines DU145, LNCaP, and PC3 and their EVs. To test the deconvolution methods’ performance on samples with EVs from cell types missing in the signature matrix, we used EVs RNA sequencing data from co-cultures of adipocytes and macrophages (Yang et al. 2017). The EV isolation procedures, characterization methods, and RNA library preparations are described in Table 2. The paired-end cell and EV RNA reads were downloaded from the NCBI GEO database (Edgar et al. 2002). Gene expression in cell and EV RNA samples was quantified at the transcript level using Salmon v1.10.2 (Patro et al. 2017) to the human Gencode v44 transcriptome. The transcript quantifications were summarized to gene level with the tximeta package (Love et al. 2020) in R version 4.3.2.

**Table 2:** Samples used for benchmarking the deconvolution methods.

| Cells/<br>organisms | EV<sup>1</sup><br>source | EV<br>isolation<sup>2</sup> | RNA<br>library kit | EV<br>validat<br>ion<sup>3</sup> | GEO/Data<br>source | Reference |
| --- | --- | --- | --- | --- | --- | --- |
| CRC | CCM | UF/dUC | NEBNext | EM<sup>4</sup> | GSE121964 | (Hinger et al. 2018) |
|  |  |  |  | WB<sup>4</sup> |  |  |
| **DU145**<br>**LNCaP**<br>**PC3** | CCM | dUC | SMARTer | NTA<br>EM<br>WB | GSE183070 | (Almeida et al. 2022) |
| **Adipocytes**<br>**Macrophages** | CCM | dUC | SMART-Seq v4 | WB | GSE94155 | (Yang et al. 2017) |
| **Human** | Plasma | exoRNeasy | SMARTer | EM<br>nanoFCM<br>WB | [exorbase.org](https://www.exorbase.org) | (Li et al. 2019; Yu et al. 2020) |
| **Human** | Urine | exoRNeasy | SMARTer | - | [exorbase.org](https://www.exorbase.org) | (Lai et al. 2021) |
| **Human** | Urine | dUC | SMART-Seq | NTA<br>EM<br>WB | Supplement Table S9 of the paper. | (Dwivedi et al. 2023) |

<sup>1</sup> CCM: cell-conditioned medium.
<sup>2</sup> UF: ultrafiltration; dUC: differential ultracentrifugation, exoRNeasy: a kit that isolates EVs by their affinity to a proprietary membrane.
<sup>3</sup> EM: electron microscopy; NTA: nanotracking analysis; nanoFCM: nano-flow cytometry; WB: western blot.
<sup>4</sup> Validated in Higginbotham et al. (2011) and Heckler et al.(2013)

### Tissue, plasma EVs, and urine EVs RNA data

We used RNA sequencing data from 27 tissues, which was downloaded from the GTEx Portal as reads (GTEx_Analysis_2017-06-05_v8_RNASeQCv1.1.9_gene_reads) and transcripts per million (GTEx_Analysis_2017-06-05_v8_RNASeQCv1.1.9_gene_tpm). We included RNA sequencing data from patients with a death classification below 4 on the HARDY Scale (i.e., excluding samples from patients that died of long-term illness). CIBERSORTx has a file size limit of 1 GB, and we, therefore, included a maximum of 200 consecutive samples from each tissue. The number of samples included for each tissue is shown in Supplementary Table S1. Plasma EV RNA sequencing data was obtained from 118 healthy humans (Table 2). We used urine EV RNA sequencing data from two cohorts that differed in the procedure for EV isolation (Table 2).

### In silico EV mixtures

To create EV samples *in silico* with precise control of mixing ratios and equal library sizes, we adapted a method for simulating RNA sequencing data from mixtures of cell types described and verified by Dong et al. (Dong et al. 2023). Briefly, this method enables the creation of *in silico* mixtures by combining randomly selected reads from pure RNA sequencing samples. To verify the approach, we re-analyzed RNA sequencing data from pure and mixed samples of NCI-H1975 and HCC827 cells (Holik et al. 2017, data accessible at NCBI GEO database, accession GSE64098) similar to Dong et al. (Dong et al. 2023). The samples consist of RNA sequencing data from H1975 and HCC827 cells mixed physically at five different ratios: 0:100, 25:75, 50:50, 75:25, 100:0. *In silico* mixtures where created by randomly extracting 7.5 million, 15 million, and 22.5 million reads from each pure H1975 and HCC827 sample using seqtk (https://github.com/lh3/seqtk) with a fixed random seed number to keep the pairing of the paired-end reads. The downsampled H1975 sequencing files were merged with downsampled HCC827 sequencing files to create an *in silico* mix with 30 million reads in ratios of 75:25, 50:50, and 25:75 in 3 replicates. Moreover, we also made mixtures of pure H1975 and HCC827 cells *in silico* by combining two 15 million read subsamples for each cell type. To make *in silico* mixed EV samples, the DU145 and LNCaP-derived EV RNA sequencing files were downsampled and mixed to create *in silico* mixed files with 30 million reads containing DU145 and LNCaP EVs in 75:25, 50:50, and 25:75 ratios in 3 replicates. Moreover, *in silico* mixtures of pure DU145 and LNCaP EV samples were also created in triplicate by combining two downsampled 15 million read files. We used a similar approach to mix pure DU145 EV samples with the EV RNA from co-cultured adipocytes and macrophages.

Count matrices were produced as described above.

### Transcriptional deconvolution methods

We used 11 deconvolution methods (Table 1) that use different algorithms to identify signature genes and the subsequent regression to estimate cell type proportions. The data were used as raw counts or counts per million (CPM) normalized as specified below.

Bisque (Jew et al. 2020) was run using the Bisque Bioconductor R package with default setting and tissue/cell and EV RNA as count matrices.

CIBERSORTx (Newman et al. 2019) was run on the website https://cibersortx.stanford.edu in absolute mode with B-mode batch correction and without quantile normalization. Tissue/cell and EV RNA expression data were CPM-normalized.

DeconRNASeq (Gong and Szustakowski 2013) was run using the DeconRNASeq v1.26 Bioconductor R package with default parameters. Tissue/cell and EV RNA expression data were CPM-normalized.

dtangle (Hunt et al. 2018) was run in R using the default setting. Tissue/cell and EV RNA expression data were CPM-normalized.

EV-origin (Li et al. 2020) was run using the R script downloaded from https://github.com/HuangLab-Fudan/EV-origin/blob/master/EV-origin.new.txt. The EV-origin script was run with default settings. Cell and EV RNA expression data were CPM-normalized. In contrast to Li et al. (2020), who used a limited list of genes with high tissue/cell type-specificity, we included the 10.000 and 20.000 genes with the highest expression variability between GTEX tissue and cells, respectively, to better compare the performance of EV-origin to the other deconvolution methods.

DWLS (Tsoucas et al. 2019) was run using the default setting in R. Tissue/cell RNA data were used as count matrices, and EV data as CPM normalized data. The function solveDampenedWLS() was used to produce the DWLS estimates. The DWLSj estimates were made by the functions findDampeningConstant() and then solveDampenedWLSj(). The DWLS OLS and DWLS SVR estimates were produced with the functions solveOLS() and solveSVR().

MuSiC v0.1.1 (Wang et al. 2019) was run using the music_prop() function from the R package available at https://github.com/xuranw/MuSiC. The estimates of the MuSiC and MuSiC nnls algorithms were retrieved from the music_prop() function output data frames Est.prop.weighted and Est.prop.allgene. Raw count data was used as input for both signatures and mixtures.

### Benchmarking scores

The deconvolution methods’ estimates were converted to relative proportions, and the total sum within one sample is always 1. Thus, a change in the abundance of one tissue’s/cell type’s EV affects the estimate of other tissue’s/cell type’s EV abundance. To assess the overall sample-to-sample performance by the deconvolution methods, we used the Bray-Curtis dissimilarity metric. This metric uses the deconvolution methods proportion estimates as a multidimensional data point, e.g., a sample consisting of EVs from the four cell lines DU145, LNCaP, PC3, and CRC is a four-dimensional data point. The accuracy of the deconvolution methods is how similar the estimates of multidimensional data points are to the true data points. The Bray-Curtis dissimilarity was calculated as:

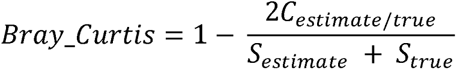

Where *C_estimate/true_* is the sum of the lesser values for each cell type EVs in estimates and true samples, *S_estimate_* and *S_true_* are the total number of EVs in the estimate and true samples. For visualization purposes, we used the Bray-Curtis index, defined as 1-Bray-Curtis dissimilarity. Thus, a Bray-Curtis dissimilarity value and Bray-Curtis index of 0 and 1, respectively, indicate that the proportions in the estimate and true sample are similar. Moreover, the root mean squared error (RMSE) was used to measure the standard deviation of the estimation error, and Pearson’s correlation coefficient *R* was used to measure the correlation between estimated and true cell type EV abundances.

### Data presentation

Sample numbers, definitions of values, and error bars are described in the figure legends. The analyses and data visualizations were done with R version 4.3.2.

## Results

### Deconvolution of EVs from cultured cells

For our benchmarking strategy, we used RNA sequencing data from four different cell lines and their EVs isolated from cell-conditioned medium (Figure 1a). A multidimensional scaling plot of the cells and EVs showed cell type-specific clustering (Figure 1b), and we employed the 11 deconvolution methods to test their ability to estimate cell type-specific EV abundances in these pure samples (Figure 1a). Compared to the truth (Figure 1c, top left, Supplementary Figure S1), the four deconvolution methods, i.e., DWLS, DWLSj, CIBERSORTx, and DWLS OLS, created Bray-Curtis index scores > 0.75 with the DWLS method estimates closest to the true samples (Bray-Curtis index 0.92, Figure 1c and d). Thus, some of the deconvolution methods estimate cell-specific EV abundances of pure samples with high accuracy.

**Figure 1:**
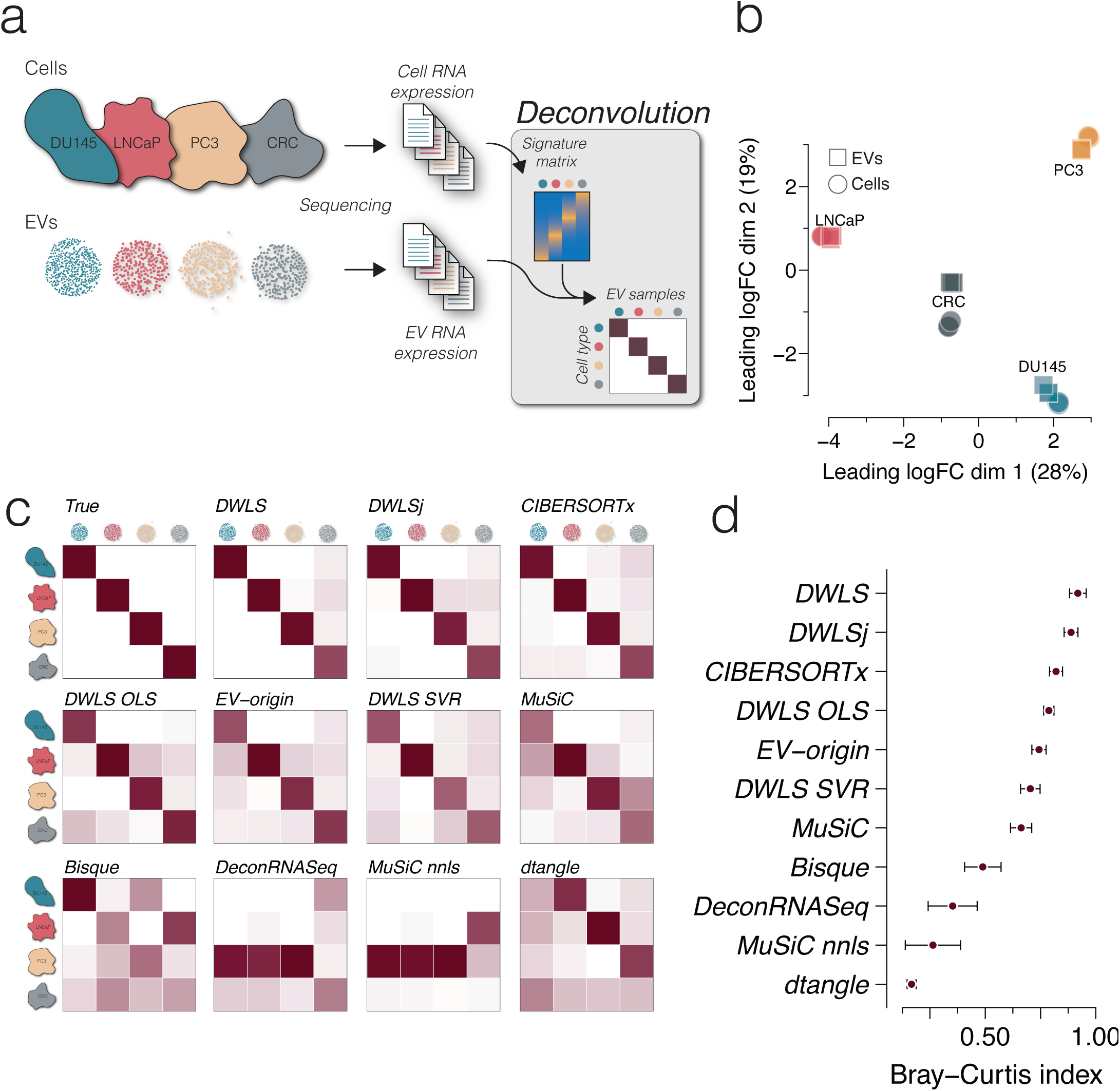
Deconvolution accuracy across methods in pure EV samples from cell cultures. **(a)** Cell and EV RNA sequencing data from 3 prostate cancer cell lines DU145, LNCaP, and PC3 (GSE183070) and a colorectal cancer (CRC, GSE121964) cell line (n = 3 cell and EV sample for each cell type). The cell RNA data was used to create a signature matrix for deconvolution of the cell-derived EV RNA data. **(b)** Multidimensional scaling plot of the cell and EV samples showing cell type-specific clustering. Circles indicate cells, and squares indicate EVs (n = 3 for each sample type). **(c)** Heatmaps of deconvolution results of the EV samples. The estimated EV abundance is the average of triplicates and is scaled between 0 and 1. The estimate for each sample is shown in Supplementary Figure S1. The numbers are represented by the darkness of the color, i.e., 0 is white, and 1 is dark red. The upper left heatmap shows the true distribution. **(d)** Similarity to the true samples (upper left heatmap) is calculated as the Bray – Curtis index for all 12 samples, i.e., 4 cell types and 3 replicates. The points indicate the mean Bray-Curtis index values and error bars indicate SEM.

EVs isolated from body fluids are a mixture derived from various cell types. To evaluate the deconvolution methods’ performance in estimating cell type-specific EV proportions in mixed samples, we mixed the pure DU145 and LNCaP EV samples *in silico* (Figure 2a). The *in silico* mixing strategy was previously described and validated by Dong et al. (2023). Briefly, RNA sequencing reads were randomly extracted from pure samples, and the downsampled reads were mixed to control mixing ratios precisely and maintain equal library size (Figure 2a). This strategy was verified by re-analyzing physically mixed NCI-H1975 and HCC827 cells, demonstrating highly similar multidimensional scaling plot clustering and gene expression profiles to the *in silico* mixtures made from pure samples (Supplementary Figure S2). The *in silico* mixing strategy was used on pure DU145 and LNCaP EV samples, which were downsampled to 22.5, 15, and 7.5 million reads. The downsampling of the pure EV samples did not affect the cell type-specific clustering, gene expression profile (Supplementary Figure S3), or the estimates of cell type-specific EV abundances by the 11 deconvolution methods (Supplementary Figure S4). The downsampled DU145 and LNCaP EV samples were randomly combined in ratios of 100:0, 75:25, 50:50, 25:75, and 0:100 in triplicate with 30 million reads per sample. A multidimensional scaling plot demonstrated that pure DU145 and LNCaP EV samples were closely clustered with *in silico* mixed DU145 and LNCaP EV samples created by combining two downsampled EV samples from the same cell type, while the *in silico* DU145/ LNCaP EV mixtures clustering was dependent on the mixing ratio (Figure 2b). The 15 *in silico* DU145/LNCaP mixtures, i.e., 5 mixing ratios in triplicate, were used to evaluate the deconvolution methods’ performances. DWLS estimated the EV proportions close to the expected ratio (Bray-Curtis index = 0.97), and the correlation between estimated and *in silico* mixed DU145 and LNCaP was high (*R_DU145_* = 0.99, *R_LNCaP_* = 0.99, Figure 2c). Similarly, DWLSj, CIBERSORTx, and DWLS OLS estimated DU145 and LNCaP EV proportions close to the expected ratios (Bray-Curtis index > 0.8, *R_DU145_* > 0.9 and *R_LNCaP_* > 0.9, Figure 2d and Supplementary Figure S5), indicating that transcriptome deconvolution can accurately predict cell type-specific EV abundances in mixed samples.

**Figure 2:**
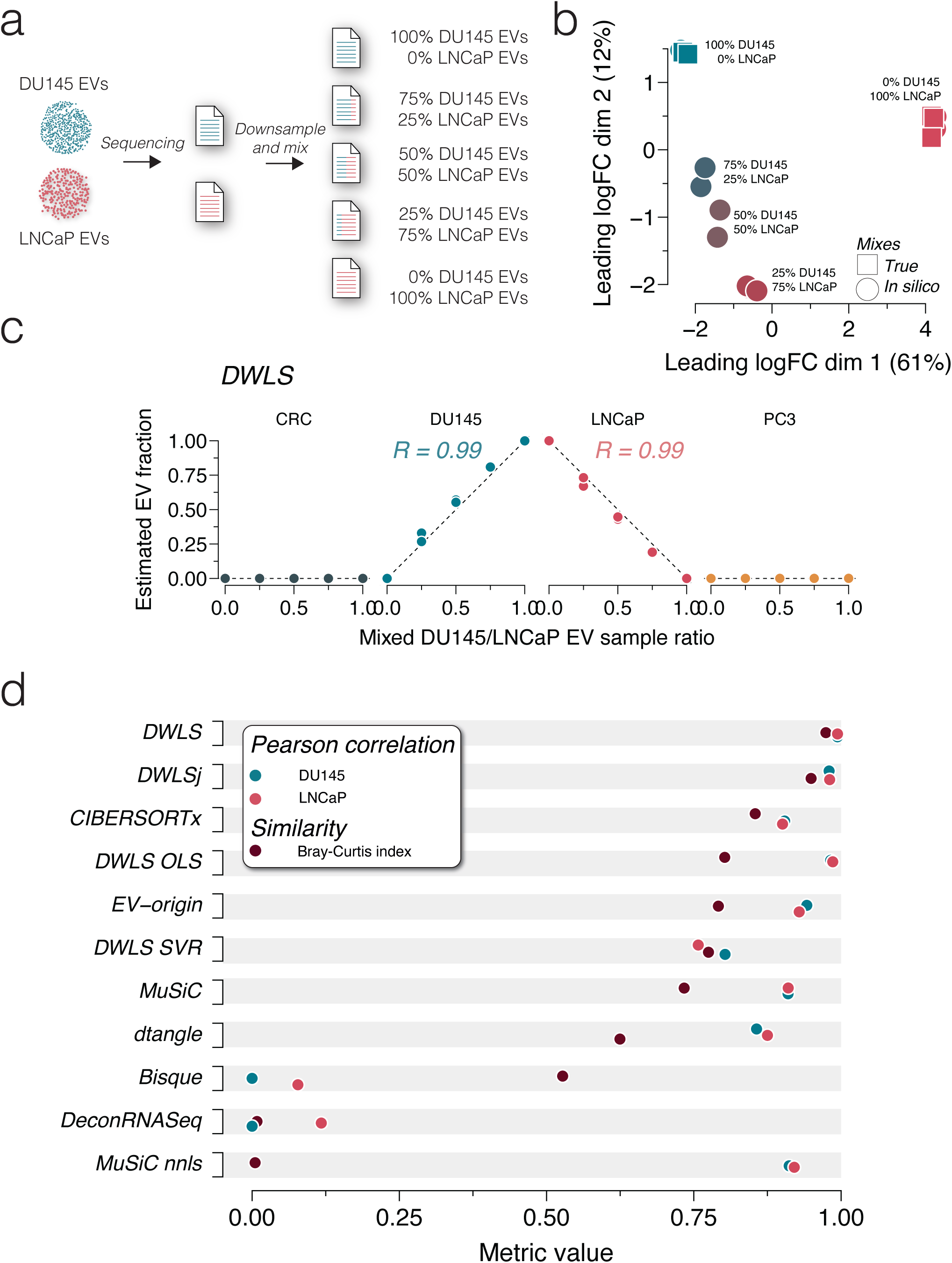
Deconvolution accuracy across methods in mixed DU145 and LNCaP EV samples. **(a)** Raw RNA sequencing reads from DU145 and LNCaP EV samples were downsampled and mixed in ratios of 100:0, 75:25, 50:50, 25:75, and 0:100. All mixed files contained 30 million reads. **(b)** Multidimensional scaling plot of true (square) and *in silico* mixed (circle) RNA sequencing reads shows ratio-specific clustering (n = 3 for each ratio). **(c)** DWLS deconvolution of DU145 and LNCaP EV samples estimates the *in silico* mixing ratio accurately. *R* indicates the Pearson correlation coefficient. The deconvolution estimates with the remaining ten methods are shown in Supplementary Figure S5. **(d)** Summary of Pearson correlation coefficients and Bray-Curtis index for deconvolution of DU145 and LNCaP *in silico* mixed EV samples for the 11 transcriptome deconvolution methods. The metrics are color-coded, as indicated in the legend.

Next, we evaluated whether EVs from cell types missing in the signature matrix interfered with the deconvolution. To this end, we made *in silico* mixed DU145 EVs with adipocyte and macrophage EV RNA (GSE94155, Yang et al. 2017) in ratios of 100:0, 75:25, 50:50, and 25:75 (Figure 3a). While samples with >50% adipocyte and macrophage EV affected the deconvolution by Bisque, DeconRNASeq, and MuSiC nnls, the EV proportions estimated by DWLS, DWLSj, CIBERSORTx, DWLS OLS, EV-origin, DWLS SVR, dtangle, and MuSiC deconvolution methods were only minimally affected by the presence of adipocyte and macrophage EV RNA even at high proportions (Figure 3b and c).

**Figure 3:**
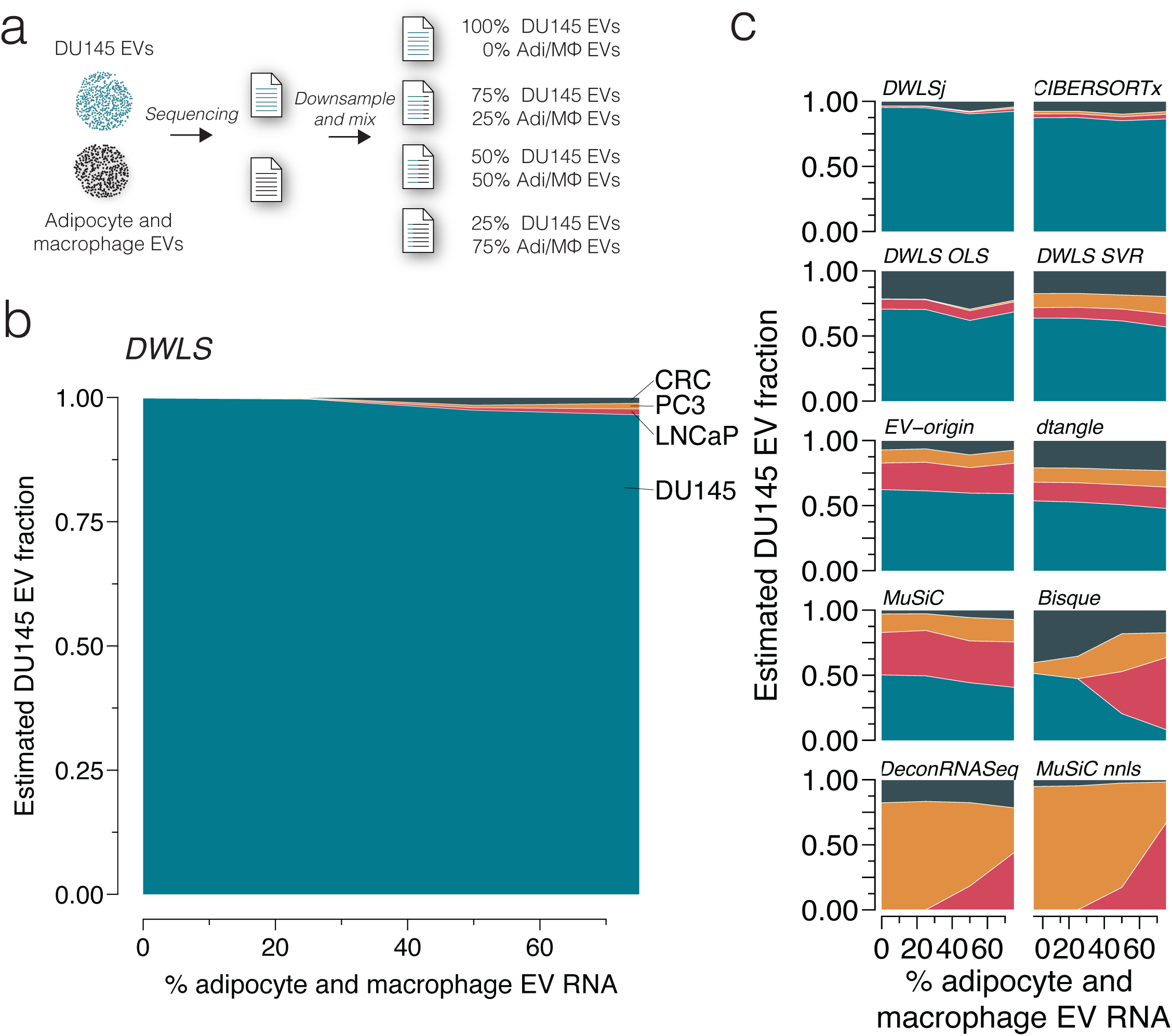
Effect of missing cell types in the signature matrix. **(a)** *In silico* mixed EV samples were created by downsampling and mixing RNA reads from DU145 and physically mixed adipocyte and macrophage EV samples (GSE94155) in ratios of 100:0, 75:25, 50:50, 25:75 (n = 3 for each ratio). The samples were deconvoluted using the transcript signatures for DU145, LNCaP, PC3, and CRC cells. **(b)** The adipocyte and macrophage EV RNA did not significantly affect the cell type-specific EV fraction estimated by DWLS. **(c)** Similarly, the cell-specific EV estimated of the mixed samples were not significantly affected by adipocyte and macrophage EV RNA for DWLSj, CIBERSORTx, DWLS OLS, DWLS SVR, EV-origin, dtangle, and MuSiC deconvolution methods; however, the presence of 50% or more adipocyte and macrophage EV RNA changed cell type-specific EV estimates for Bisque, DeconRNASeq, and MuSiC nnls deconvolution methods.

To evaluate the deconvolution methods’ performance overall, we combined their scores (RMSE, Bray-Curtis dissimilarity, and correlation coefficients) from deconvoluting pure, downsampled, and mixed samples (Figure 4). For visualization purposes, we used 1 - Pearson correlation coefficient (PCC) values in which 0 is a perfect correlation. DWLS created deconvolution results closest to the expected (Figure 4a), followed by DWLSj, CIBERSORTx, and DWLS OLS (Figure 4b). Together, our deconvolution analyses of the cell line-derived EVs demonstrate that current methods for transcriptome deconvolution enable cell type-specific EV abundance estimation with high accuracy.

**Figure 4:**
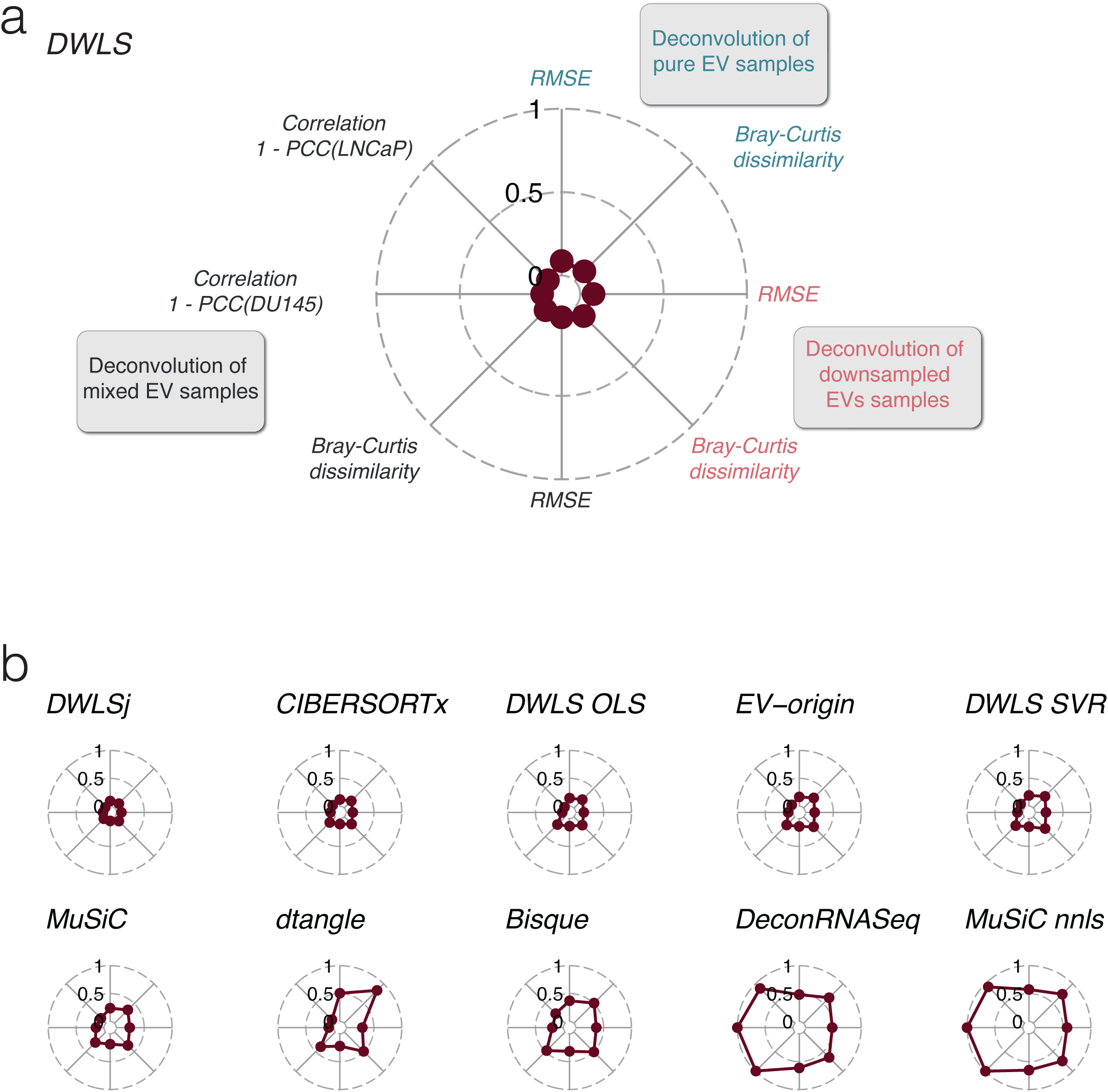
Benchmarking the deconvolution accuracy for cell line-derived EV samples across methods. **(a)** Overall benchmarking scores of DWLS deconvolution of the pure, the downsampled, and the *in silico* mixed EV samples. RMSE indicates root mean square error. Correlation for the deconvolution of the *in silico* mixed DU145 and LNCaP EV samples is shown as 1-PCC, where PCC indicates Pearson Correlation Coefficient. **(b)** Overall benchmarking of the remaining ten transcriptome deconvolution methods.

### Deconvolution of urine EVs

To evaluate the deconvolution methods’ ability to predict the tissue-specific EV abundances in urine samples, we used urine EV RNA sequencing data from two independent cohorts, which we will refer to as exoRbase (n = 16 samples, Lai et al. 2021) and Dwivedi (n = 72 samples, Dwivedi et al. 2023) cohorts. Notably, the urine EV isolation methods for the two cohorts were different: the exoRbase cohort’s urine EVs were isolated by exoRNAeasy, and the Dwivedi cohort’s urine EVs were isolated by differential ultracentrifugation.

Firstly, we created a signature matrix with tissue-specific gene expression from 27 human tissues. For the exoRbase and Dwivedi cohorts, the deconvolution methods’ tissue-specific EV abundance estimates were highly variable (Supplementary Figures S6 and S7), and the correlation coefficients between the methods’ estimates were low (Supplementary Figure S8). We noticed that epithelial-rich organs, such as the pancreas and stomach, were predicted to contribute a significant fraction of EVs to the urine. This contrasts with other lines of experimental evidence suggesting that urine EVs are predominantly derived from epithelial cells in the genitourinary system (Svenningsen et al. 2020; Blijdorp et al. 2022; Nørgård et al. 2022). We, thus, restricted the tissue signature matrix to nine tissues of the genitourinary system. Using the genitourinary-restricted tissue signature matrix for deconvolution of the exoRbase cohort samples demonstrated that the DWLS, DWLSj, CIBERSORTx, and DWLS OLS deconvolution methods estimated the kidneys to be dominant contributors of urine EVs (80 ± 6%, Figure 5a). The correlations between the DWLS, DWLSj, CIBERSORTx, and DWLS OLS deconvolution methods were high (*R* > 0.95, Figure 5b). Urine EV samples from the Dwivedi cohort were estimated by DWLS, DWLSj, CIBERSORTx, and DWLS OLS deconvolution to contain EVs derived from kidneys (49 ± 5%) and bladder (39 ± 2%), accounting for 88 ± 6% of the urine EVs (Figure 5c) and with high correlation between the four deconvolution methods estimates *R >* 0.95, Figure 5d). We then compared the deconvolution methods’ estimates for the two cohorts. The DWLS method’s deconvolution of the two cohorts was highly correlated (*R* = 0.95, Figure 5e). CIBERSORTx demonstrated a lower correlation (*R* = 0.56), but DWLSj and DWLS OLS showed high correlations between the two cohorts’ estimates (*R_DWLSj_* = 0.85 and *R_DWLS OLS_* = 0.85, Figure 5f). Although the tissue gene expression matrix is crucial for obtaining plausible results, there is a high agreement between the results created by DWLS, DWLSj, CIBERSORTx, and DWLS OLS, and there is a strong positive correlation between these deconvolution methods’ estimates between cohorts.

**Figure 5:**
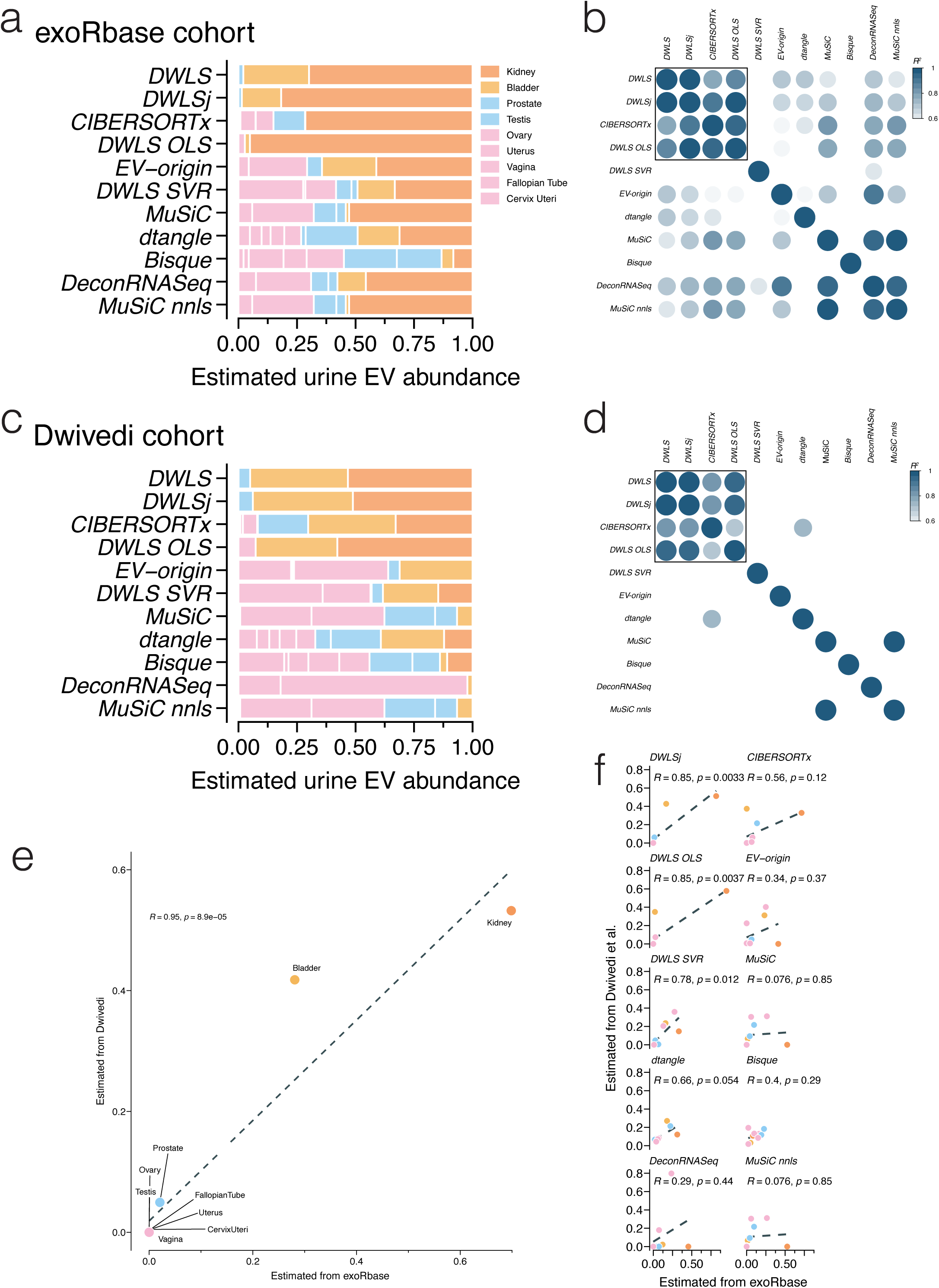
Deconvoluting of tissue-specific EV abundances in urine across methods. **(a)** Deconvolution of urine EV samples from the exoRbase cohort (n = 16 samples) by the 11 deconvolution methods. The deconvolution methods’ estimated tissue/organ source is color-coded according to the legend. **(b)** Pearson correlation coefficients (*R^2^* indicated by color) between the 11 deconvolution methods’ estimates of tissue-specific urine EV abundance. **(c)** Estimated tissue-specific EV abundance and **(d)** Pearson correlation coefficients (*R^2^* indicated by color) between the 11 deconvolution methods’ estimates of the urine EVs from the Dwivedi et al. (2023) cohort (n = 72). **(e)** Correlation between the tissue-specific urine EV abundance estimates between the two cohorts by DWLS, and **(f)** the remaining ten transcriptome deconvolution methods. R indicates the Pearson correlation coefficient.

### Deconvolution of plasma EVs

For plasma EVs, we used the 27-tissue signature to estimate the tissue-specific EV abundances for plasma samples from 118 healthy persons. While DWLS, DWLSj, and CIBERSORTx estimated blood and spleen contributed 95 ± 2% of the plasma EVs (Figure 6a), they estimated different proportions of these two tissues. CIBERSORTx estimated that 50% of the plasma EVs were derived from the spleen, but DWLS and DWLSj estimated that the spleen-derived EVs only contributed with <5% of plasma EVs (Figure 6a). Consistent with this, the correlation between the tissue-specific EV abundances estimated by the different deconvolution methods was low (Supplementary Figure S9). The spleen is a reservoir for blood cells, and we, therefore, combined the estimates for blood and spleen-derived plasma EVs. This improved the correlations between DWLS, DWLSj, CIBERSORTx, and DWLS OLS algorithms (*R* > 0.98, Figure 6b) and produced highly similar estimates for each of the 118 samples (Figure 6c). Thus, the DWLS-based and CIBERSORTx deconvolution methods had highly concordant estimates of tissue-specific EV abundances in plasma samples.

**Figure 6:**
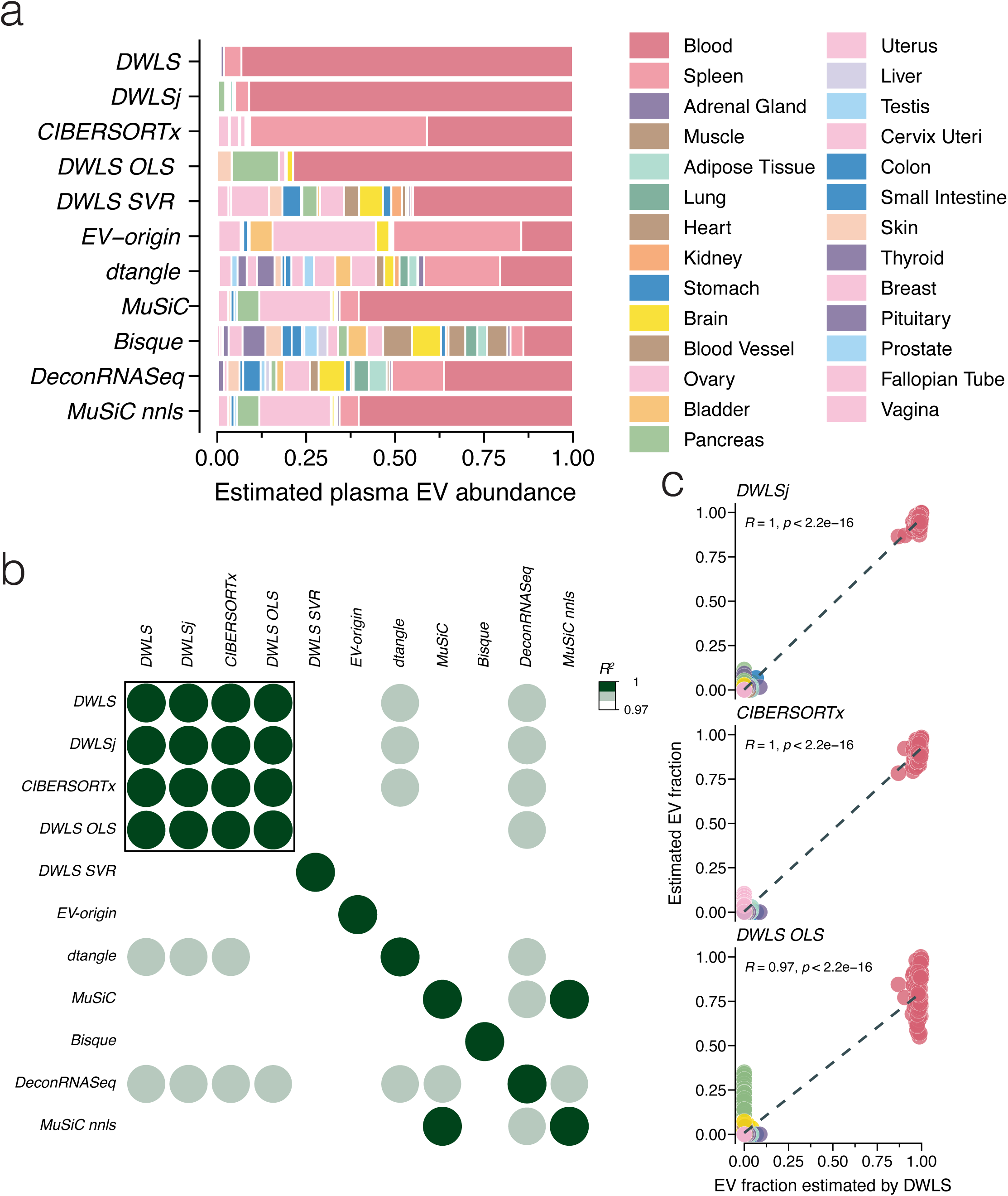
Deconvolution of tissue-specific EV abundances in plasma across methods. **(a)** Estimated tissue-specific plasma EV abundance of plasma EV samples (n = 118, GEO accession numbers GSE100206 and GSE133684) by the 11 deconvolution methods. The tissues/organs are color-coded according to the legend. **(b)** Pearson correlation coefficients (*R^2^* indicated by color) between the 11 deconvolution methods’ estimates of the tissue-specific EV abundance in plasma samples. **(c)** Correlation between the individual person’s tissue-specific plasma EV abundance estimated by DLWS and (top panel) DWLSj, (middle panel) CIBERSORTx, and (bottom panel) DWLS OLS. *R* indicates the Pearson correlation coefficient.

## Discussion

By meticulous analyses of transcriptomic deconvolution methods on EVs from cultured cells, plasma, and urine samples, we here provide comprehensive benchmarking data that can help address the question of tissue- and cell type-specificity of EVs isolated from biological fluids. We analyzed 11 deconvolution methods. By comparing their performances on several sample types, we identify four methods with highly concordant results in all the benchmarking tests. Importantly, our analyses highlighted the tissue/cell type signature matrix’s essential role in the EV abundance estimates. Overall, the benchmarking suggests that the best performing transcriptomic deconvolution methods, i.e., DWLS and CIBERSORTx, can convert EV RNA sequencing data into reliable tissue- and cell type-specific EV abundance estimates in biofluids.

The deconvolution methods differ in the algorithms used to identify the most informative signature genes, how these genes are weighted, and the final regression analyses. Three deconvolution methods, CIBERSORTx, EV-origin, and DWLS SVR, are based on support vector regression (SVR). Although the SVR-based methods’ estimates for cell line-derived EV were similar, they demonstrated low correlations in urine and plasma EV samples. CIBERSORTx’s estimates were most consistent with the best-performing deconvolution methods, DWLS, DWLSj, and DWLS OLS. While these three methods use the same algorithm to establish the tissue/cell type-specific genes by differential expression analyses, they treat this information differently. DWLS OLS uses ordinary least square regression (OLS); however, this algorithm tends to ignore low abundant transcripts, such as genes only expressed in rare cells or genes only expressed at low levels (Tsoucas et al. 2019). DWLS and DWLSj are also based on OLS regression, but they reduce this bias by a weighted least square approach to include the contribution for each gene (Tsoucas et al. 2019). Importantly, DWLSj estimates the weights based on the initial OLS regression, but DWLS uses an iterative approach that updates the weighted least square solution until the convergence of its estimates (Tsoucas et al. 2019). Consistent with these additional optimization steps, DWLS makes the most accurate estimates on the cell line-derived EVs and demonstrates the importance of identifying the most informative transcripts for deconvolution.

A significant challenge for the deconvolution methods is cell types with similar gene expression. Consistent with previous results using EV-origin and a carefully constructed tissue signature matrix (Li et al. 2020), the estimates of the four deconvolution methods showed that blood and spleen cells contribute significantly to the plasma EV pool; however, the same 27-tissue signature estimated a high abundance of urine EVs derived from epithelial-rich tissue, such as the pancreas and stomach. These initial urine EV estimates are not supported by other sources of experimental evidence (Svenningsen et al. 2020; Blijdorp et al. 2022; Nørgård et al. 2022). Yet, by restricting the number of tissues to only include tissues of the genitourinary systems in the signature matrix, the four best-performing deconvolution methods produced highly concordant estimates for the two independent urine EV cohorts that, among others, differed in their EV isolation procedure. The four best-performing deconvolution methods indicated that the bladder and kidneys are the dominant sources of urine EVs. Using CIBERSORTx, Zhu et al. (2021) have previously reported that bladder- and lung-derived EVs were abundantly present in urine, but kidney-derived EVs were rarer. The discrepancy between Zhu et al. (2021) and our findings is multifactorial, but choices of which tissues and genes to include in the signature are likely significant drivers. Nonetheless, data from transgenic EV reporter mice suggest that the kidneys do not freely filter the plasma EVs (Nørgård et al. 2022), and proteome data of human EVs sampled from a nephrostomy drain or normally voided urine support a substantial role of the kidneys as a source for urine EVs (Blijdorp et al. 2022). Thus, by including this biological knowledge of which tissues/cell types are likely to contribute, EVs can improve the deconvolution methods’ estimates.

It is important to highlight that plasma and urine EVs are not only derived from a few sources, but that many tissues/cell types contribute. Identifying these tissues/cell types is challenging with the marked-based deconvolution methods. As discussed above, too many tissues/cell types with similar gene expression may confound the results; however, the opposite is also true. Our test with the addition of EV RNA from adipocytes and macrophages, which was not present in the signature matrix, did not affect the deconvolution estimates of best-performing methods. While marker-based deconvolution methods cannot classify EVs from the “unknown” cell types, complete deconvolution methods can unmix samples without knowledge about cell types (Zaitsev et al. 2019). Nonetheless, our analyses showed that the relative abundances of the known cell type-specific EVs were maintained, demonstrating that the deconvolution estimates are highly sensitive to the composition of the signature matrix.

Our benchmarking indicates that the transcriptome deconvolution of EV RNA data from biofluid is an attractive means to unmix tissue/cell type-specific EV abundances and thereby provide an additional method to aid the interpretation of bulk analyses of EV cargo such as proteins, lipids, and RNAs. EV cargo abundance is a product of two main factors: the level of cargo within each EV and the abundance of cell type-specific EVs in the sample. The relative contribution of these two factors is dynamically regulated by cellular gene expression level and cell type-specific EV release rate. While isolation or enrichment for cell type-specific EVs could unmix these two factors, only a restricted number of affinity reagents are currently available. This constraint does not limit the deconvolution methods, which can effectively quantify the influence of the tissue/cell type-specific EV abundance.

Our overall recommendations for transcriptome deconvolution of EV samples are that i) DWLS (Tsoucas et al. 2019) and CIBERSORTx (Newman et al. 2019) are good method choices, ii) cellular subtypes or tissues with highly similar cell type composition can significantly bias the EV estimates, and iii) the tissue/cell type signature matrix should match biological knowledge and cover the tissue/cell types expected to contribute to the biofluid’s EV pool. Although additional optimization on the genes included in the signature matrix may improve the estimates of the other methods, DWLS and CIBERSORTx made accurate estimates using their default settings for the construction of the signature matrix. Moreover, their highly similar and robust estimates of the DWLS and CIBERSORTx methods suggest that these two existing transcriptome deconvolution methods offer tools to determine tissue- and cell type-specific EV abundances. Combined with expression analyses and, e.g., proteomics data, this can provide new insights into the role of EVs in health and diseases.

## Acknowledgments

This work was supported by the Independent Research Fund Denmark under Grant #10.46540/3103-00263B; and the Augustinus Fonden under Grant #22-2015. Alfred Gandrup Svenningsen is thanked for helping to download the data from the GEO database and transcript quantification.

---

**Supplementary Table S1:** Number of tissue samples included from the GTEX portal.

| **Tissue** | **Samples (n)** |
| --- | --- |
| Adipose Tissue | 200 |
| Adrenal Gland | 94 |
| Bladder | 14 |
| Blood | 200 |
| Blood Vessel | 200 |
| Brain | 200 |
| Breast | 120 |
| Cervix Uteri | 17 |
| Colon | 193 |
| Fallopian Tube | 8 |
| Heart | 200 |
| Kidney | 22 |
| Liver | 62 |
| Lung | 169 |
| Muscle | 200 |
| Ovary | 58 |
| Pancreas | 122 |
| Pituitary | 26 |
| Prostate | 68 |
| Skin | 200 |
| Small Intestine | 60 |
| Spleen | 81 |
| Stomach | 131 |
| Testis | 109 |
| Thyroid | 173 |
| Uterus | 50 |
| Vagina | 54 |

