## Supplementary figures and images for "Benchmarking transcriptional deconvolution methods for estimating tissue- and cell type-specific extracellular vesicle abundances"

### Supplementary Figure 1

#### Supplemental Figure S1

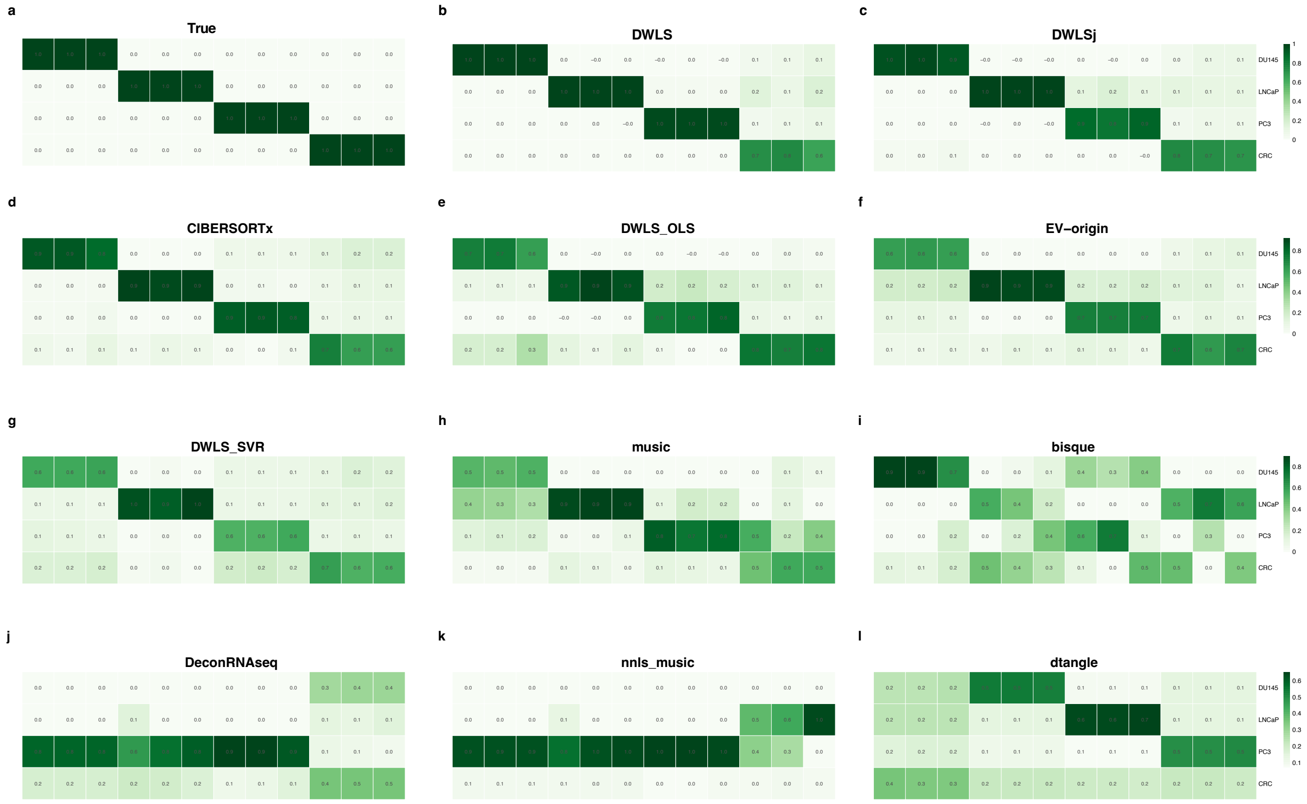

### Supplementary Figure 2

#### Supplemental Figure S2

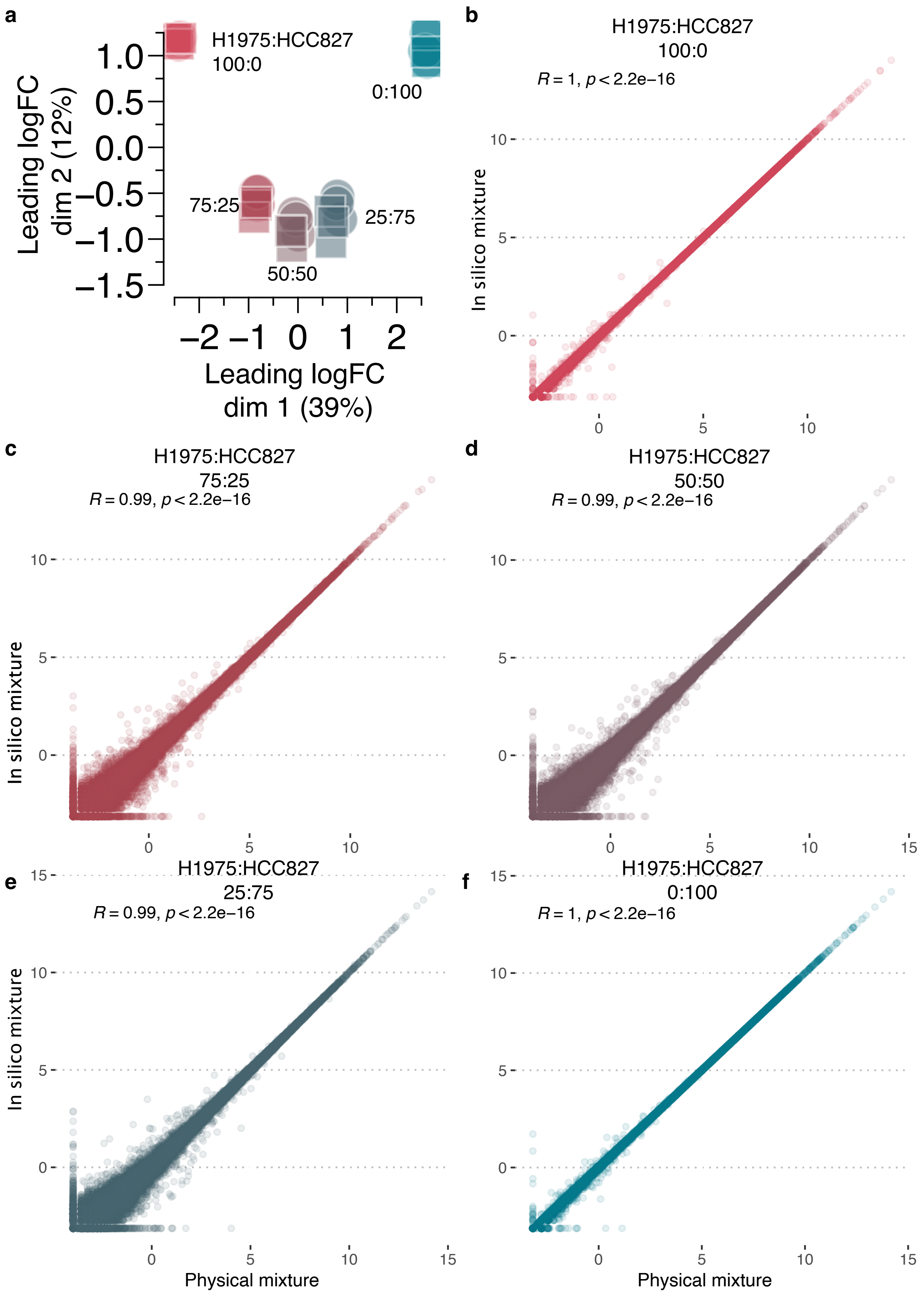

### Supplementary Figure 3

#### Supplemental Figure S3

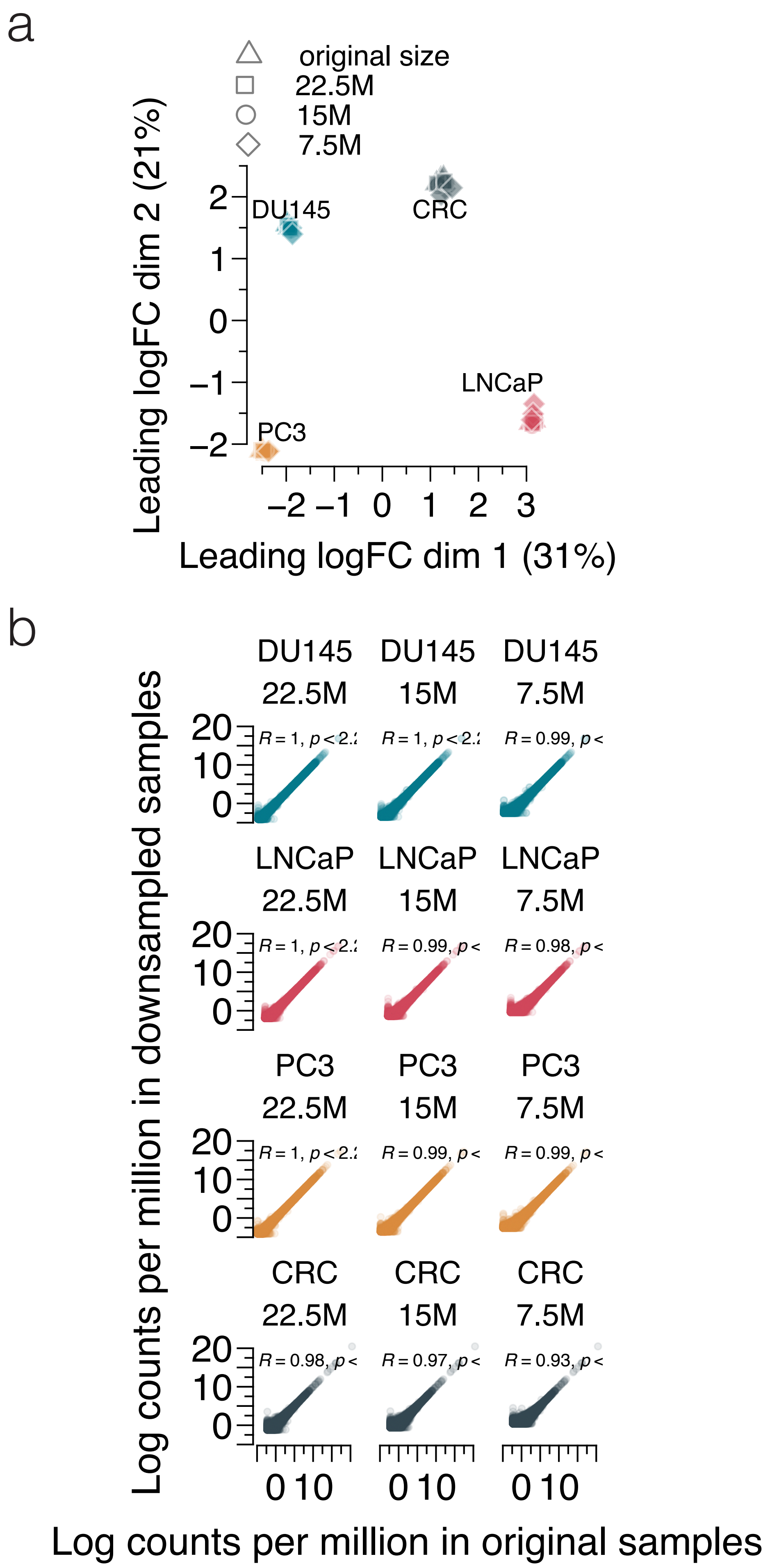

### Supplementary Figure 4

#### Supplemental Figure S4

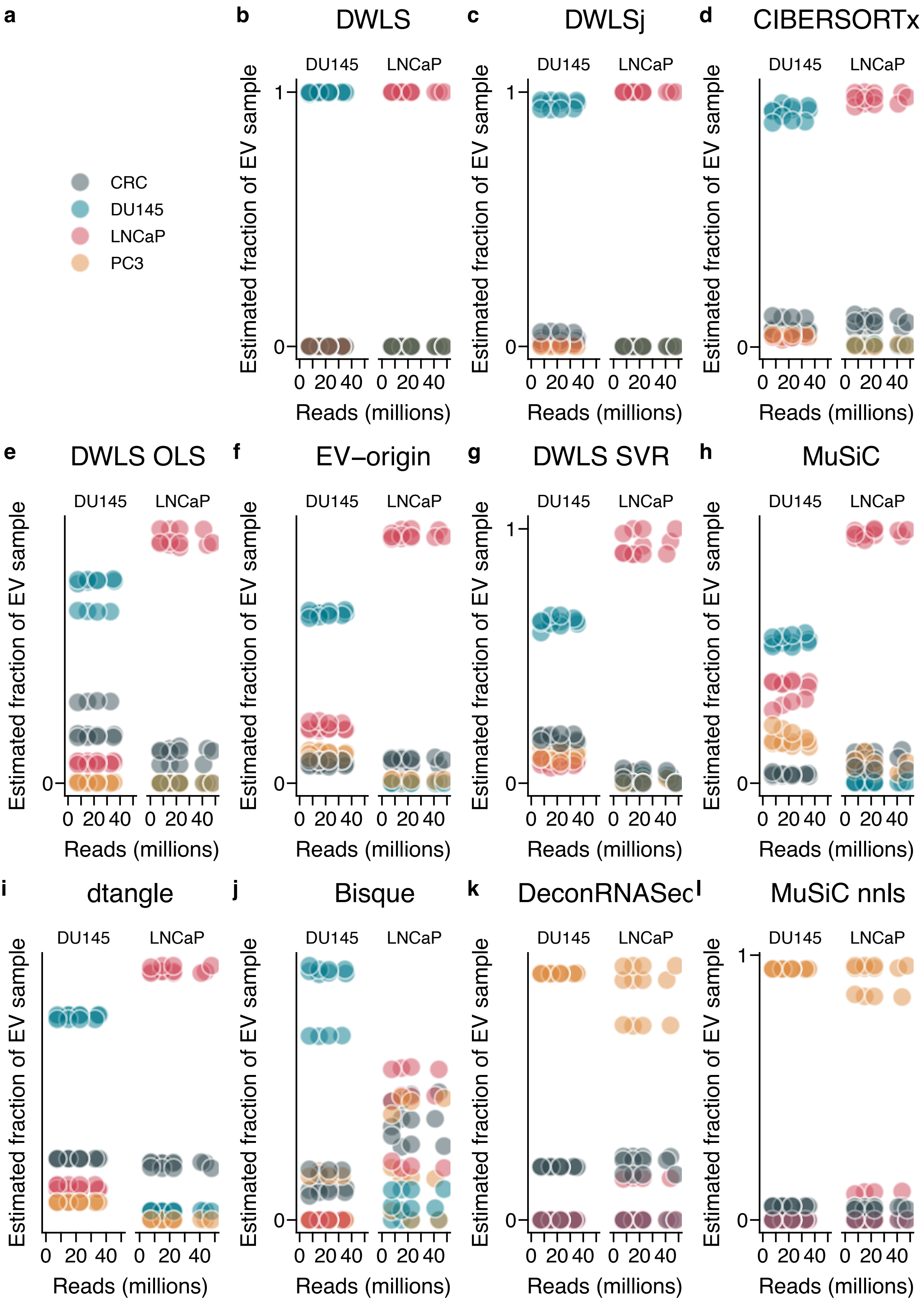

### Supplementary Figure 5

#### Supplemental Figure S5

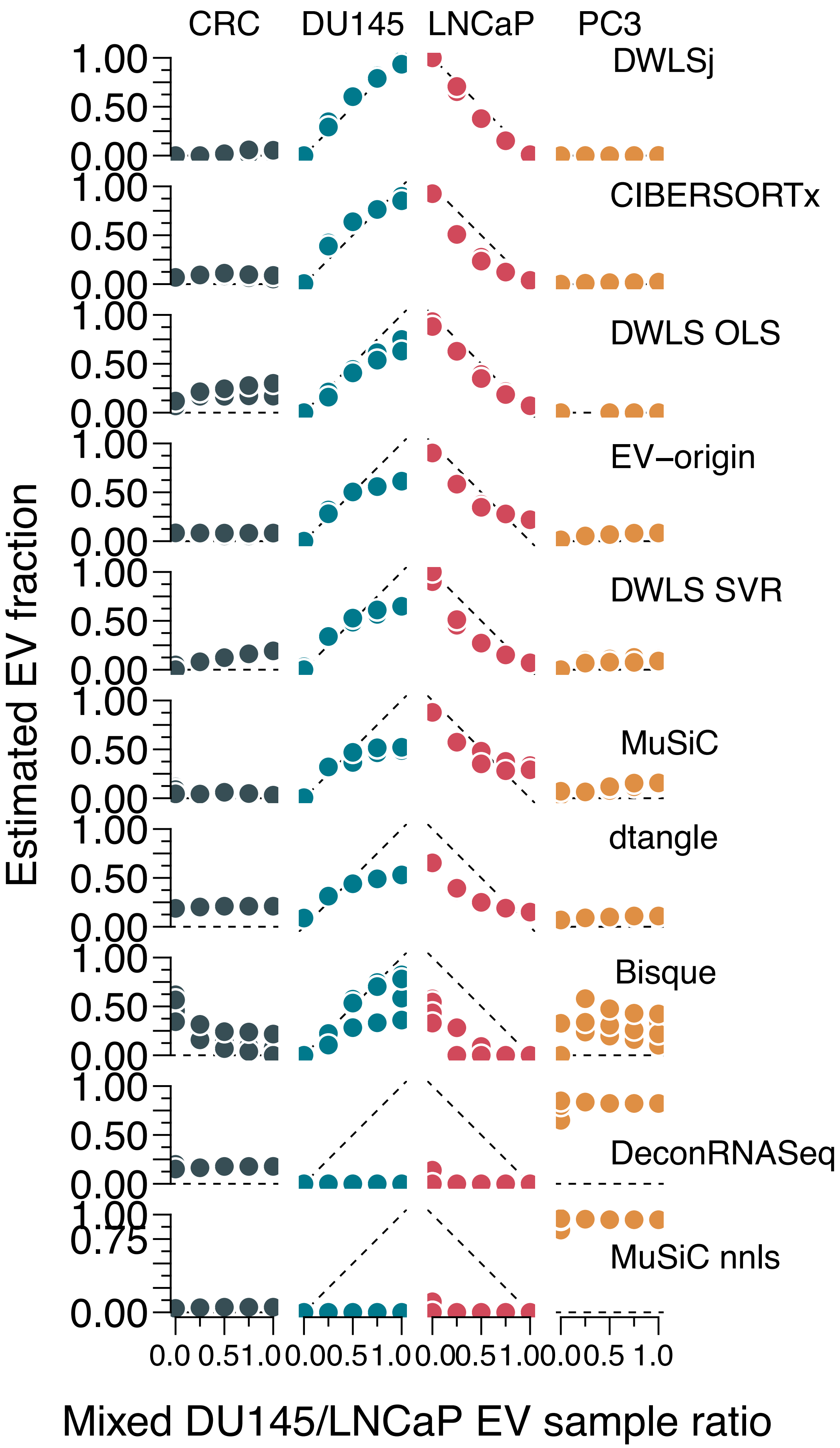

### Supplementary Figure 6

#### Supplemental Figure S6

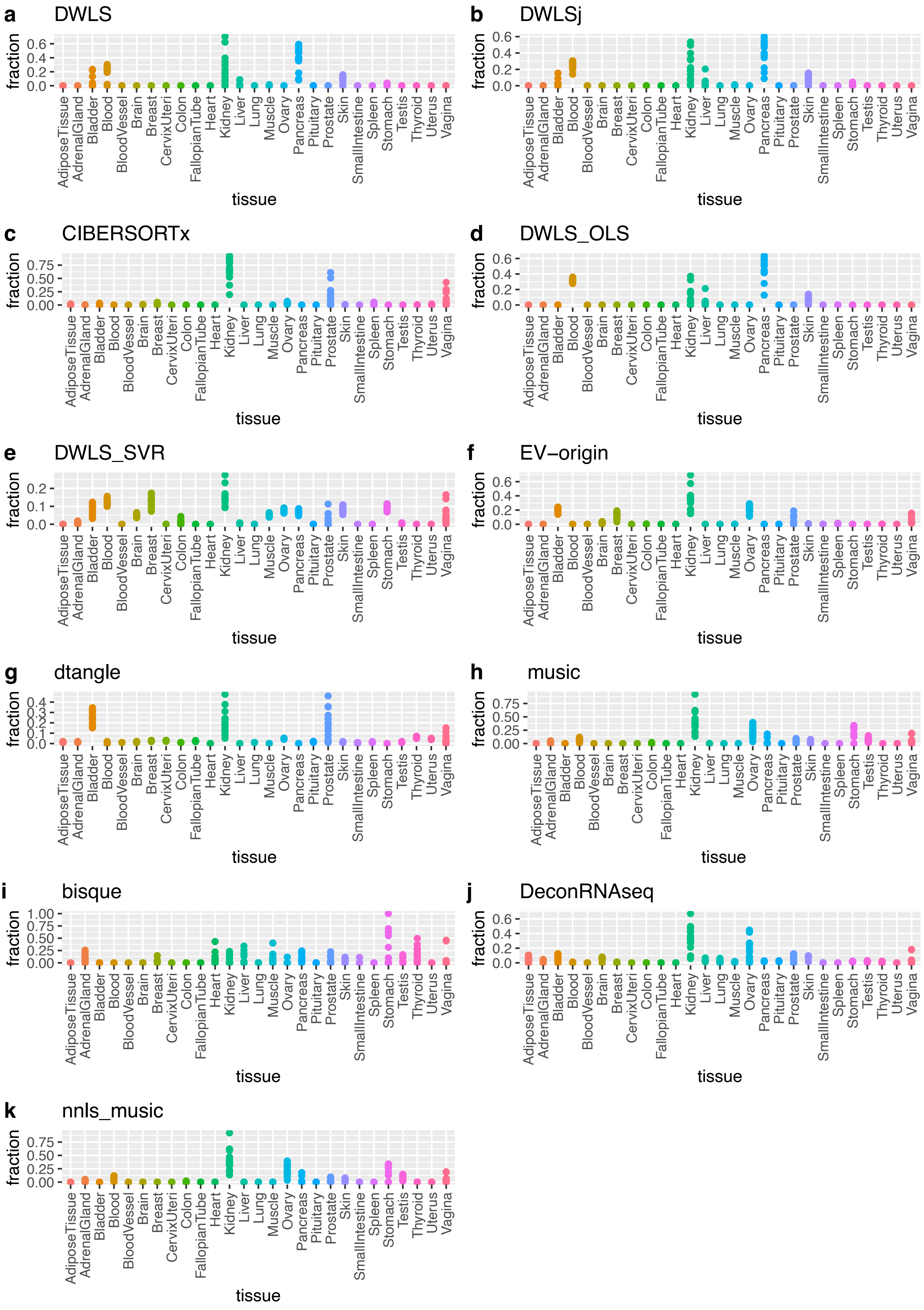

### Supplementary Figure 7

#### Supplemental Figure S7

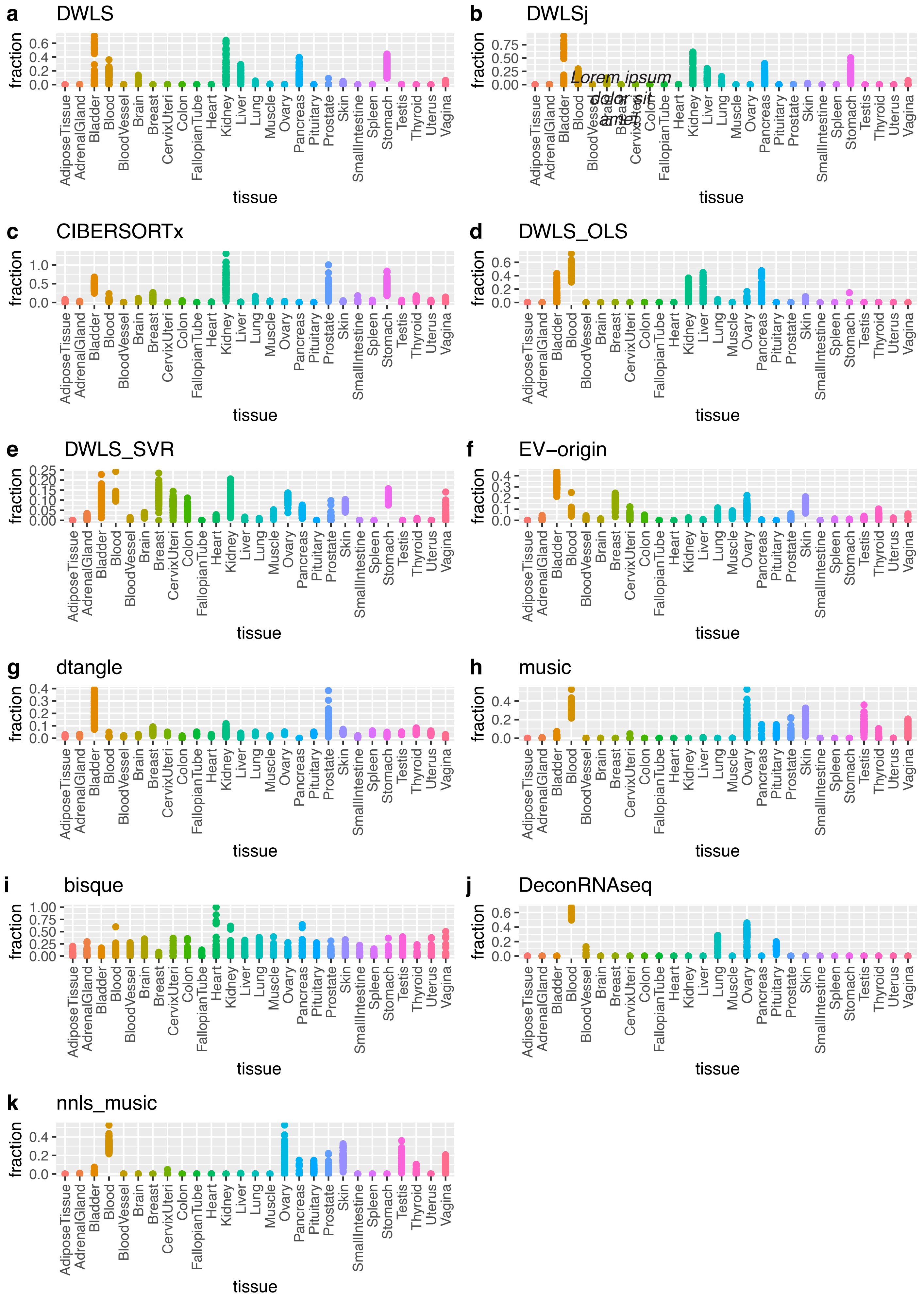

### Supplementary Figure 9

#### Supplemental Figure S9

Deconvolution of plasma EVs without merging blood and spleen

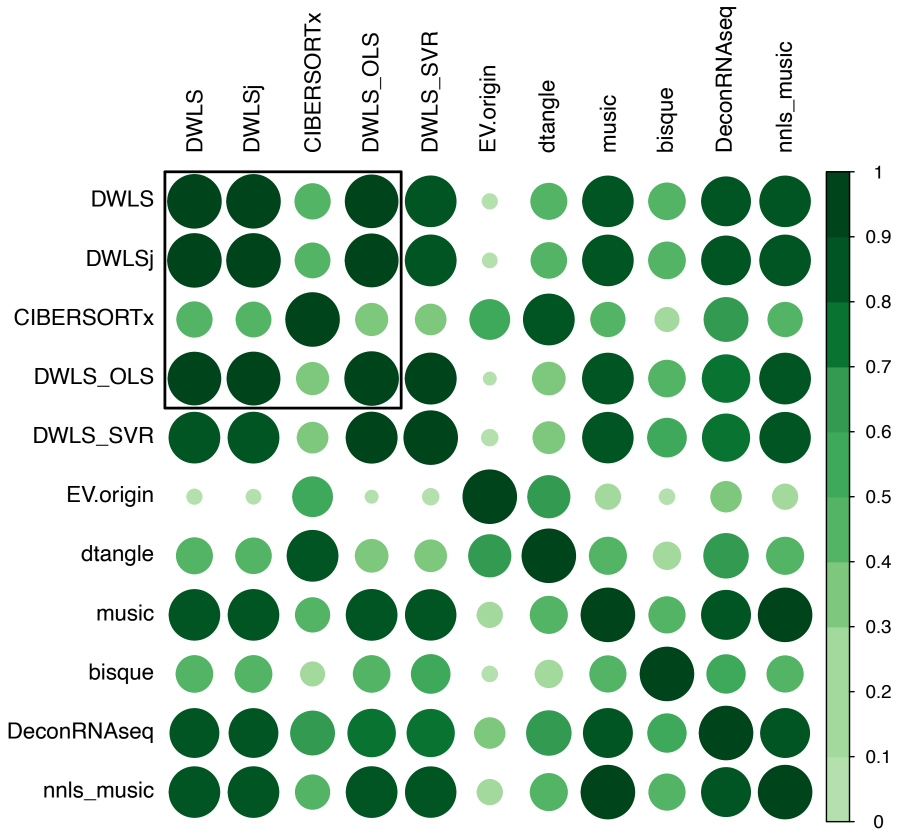
