## Supplementary Figure 8 for "Benchmarking transcriptional deconvolution methods for estimating tissue- and cell type-specific extracellular vesicle abundances"

### Supplemental Figure S8

a

Deconvolution of exoRbase corhort uEVs with 27-tissue signature matrix

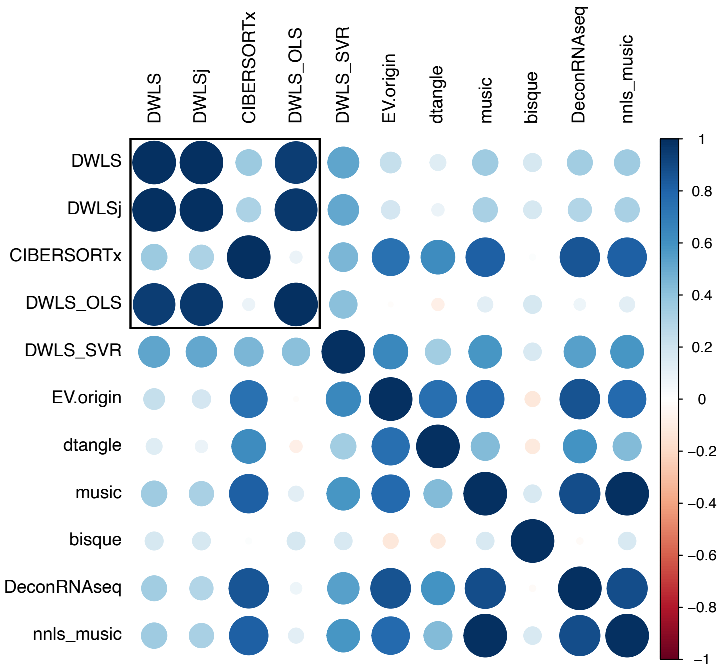


b

Deconvolution of Dwiverdi corhort uEVs with 27-tissue signature matrix

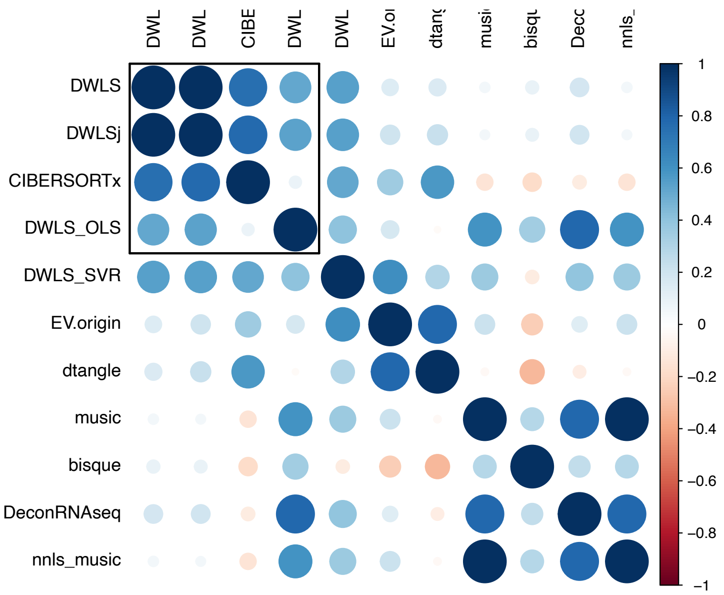
